# The Unified Phenotype Ontology (uPheno): A framework for cross-species integrative phenomics

**DOI:** 10.1101/2024.09.18.613276

**Authors:** Nicolas Matentzoglu, Susan M Bello, Ray Stefancsik, Sarah M. Alghamdi, Anna V. Anagnostopoulos, James P. Balhoff, Meghan A. Balk, Yvonne M. Bradford, Yasemin Bridges, Tiffany J. Callahan, Harry Caufield, Alayne Cuzick, Leigh C Carmody, Anita R. Caron, Vinicius de Souza, Stacia R. Engel, Petra Fey, Malcolm Fisher, Sarah Gehrke, Christian Grove, Peter Hansen, Nomi L. Harris, Midori A. Harris, Laura Harris, Arwa Ibrahim, Julius O.B. Jacobsen, Sebastian Köhler, Julie A. McMurry, Violeta Munoz-Fuentes, Monica C. Munoz-Torres, Helen Parkinson, Zoë M Pendlington, Clare Pilgrim, Sofia MC Robb, Peter N. Robinson, James Seager, Erik Segerdell, Damian Smedley, Elliot Sollis, Sabrina Toro, Nicole Vasilevsky, Valerie Wood, Melissa A. Haendel, Christopher J. Mungall, James A. McLaughlin, David Osumi-Sutherland

**Author notes:** Joint first authors (Nicolas Matentzoglu, Susan M Bello, Ray Stefancsik).

## Abstract

Phenotypic data are critical for understanding biological mechanisms and consequences of genomic variation, and are pivotal for clinical use cases such as disease diagnostics and treatment development. For over a century, vast quantities of phenotype data have been collected in many different contexts covering a variety of organisms. The emerging field of phenomics focuses on integrating and interpreting these data to inform biological hypotheses.

A major impediment in phenomics is the wide range of distinct and disconnected approaches to recording the observable characteristics of an organism. Phenotype data are collected and curated using free text, single terms or combinations of terms, using multiple vocabularies, terminologies, or ontologies. Integrating these heterogeneous and often siloed data enables the application of biological knowledge both within and across species. Existing integration efforts are typically limited to mappings between pairs of terminologies; a generic knowledge representation that captures the full range of cross-species phenomics data is much needed.

We have developed the Unified Phenotype Ontology (uPheno) framework, a community effort to provide an integration layer over domain-specific phenotype ontologies, as a single, unified, logical representation. uPheno comprises (1) a system for consistent computational definition of phenotype terms using ontology design patterns, maintained as a community library; (2) a hierarchical vocabulary of species-neutral phenotype terms under which their species-specific counterparts are grouped; and (3) mapping tables between species-specific ontologies. This harmonized representation supports use cases such as cross-species integration of genotype-phenotype associations from different organisms and cross-species informed variant prioritization.

## 1. Introduction

Phenotypes are observable or measurable characteristics of an organism resulting from the interaction of its genotype with the environment. Collecting and analyzing information about phenotypes, known as *phenotyping*, is fundamental to biological science and has many applications: clinicians record a patient’s phenotypic profile to facilitate a more accurate diagnosis; researchers record phenotypic profiles of model organisms to assess interventions (genetic or drug or otherwise); database curators integrate phenotype data with other data types by extracting phenotypes from data sources that are typically unstructured. As the body of phenotype data has expanded, researchers have looked for ways to use this collective knowledge, but the disparate methods used to record these data have posed an impediment.

Variation in alleles of orthologous genes can result in similar phenotypes across species and taxa. For example, *PAX6* gene mutations can lead to human eye phenotypes similar to the mouse phenotypes caused by *Pax6*-ortholog alleles^1^. Based on similar *FOXP2* phenotypes across humans, primates, mice and even birds, important inferences can be made about neural mechanisms that contribute to the evolution of human spoken language^2^. Evolutionarily conserved functions of myostatin gene orthologs manifest in similar muscle growth phenotypes across several vertebrate species^3^. All these examples suggest that phenotypic similarity frequently correlates with the conserved function of gene products and regulatory networks. Identifying similar phenotypes across species can provide not only supporting evidence for conserved gene function, but also the possibility of modeling phenotypes in experimentally accessible model organisms to facilitate useful discoveries in agricultural or medical research.

Phenotype ontologies have been developed to reduce ambiguity and relate similar phenotypes, making it possible for computational methods to group phenotypes easily. However, these ontologies have been developed to meet specific use cases and they are widely used in those communities. For example, the Human Phenotype Ontology (HPO)^4^ has been designed to provide a standardized vocabulary of phenotypic abnormalities and clinical features encountered in human disease and is a recognized standard for the computational encoding of human phenotyping data. HPO enables computational inference, supports genomic and phenotypic analyses, and has widespread applications in clinical diagnostics and translational research. Similarly, the Mammalian Phenotype Ontology (MP)^5^ has been developed to meet the needs of mammalian model organism data. The specific needs of each community influence the design of each ontology. As a result, despite significant overlap between species-specific ontologies, there are notable differences in their axiomatization, classification and coverage.

As the evolutionary distance between the species increases, the differences in their phenotype ontologies also expands. Ontologies for describing phenotypes of the fruit fly *Drosophila melanogaster* (Drosophila Phenotype Ontology, DPO)^6^, nematode *Caenorhabditis elegans* (*C. elegans* Phenotype Ontology, WBPhenotype)^7^ and the fission yeast *S. pombe* (Fission Yeast Phenotype Ontology, FYPO)^8^, developed to address specific needs of the respective communities, differ extensively in terms of both term organization (the taxonomy, i.e. hierarchy of terms) and the scope (for example, anatomical, cellular or molecular level) and granularity of phenotypes covered. (See Table 1 for a list of eukaryotic single and multicellular species-specific phenotype ontologies developed to describe scientific data in their respective communities).

**Table 1.** Domain-specific phenotype ontologies currently integrated into uPheno.

| Ontology | Taxon | Term Count (release version) | Reference |
| --- | --- | --- | --- |
| Mammalian Phenotype Ontology (MP) | Mammalia | 14,206 (v2024-08-08) | <sup>5</sup> |
| Human Phenotype Ontology (HPO) | *Homo sapiens* | 18,987 (v2024-08-13) | <sup>4</sup> |
| Zebrafish Phenotype Ontology (ZP) | *Danio rerio* | 47,443 (v2024-04-18) | https://github.com/obophenotype/zebrafish-phenotype-ontology |
| C. elegans Phenotype Ontology (WBPhenotype) | Nematoda | 2,649 (v2024-06-05) | <sup>7</sup> |
| Drosophila Phenotype Ontology (DPO) | Drosophilidae | 253 (v2024-04-25) | <sup>6</sup> |
| Dictyostelium Phenotype Ontology (DDPHENO) | *Dictyostelium discoideum* | 1,017 (v2023-08-26) | <sup>29</sup> |
| Planarian Phenotype Ontology (PLANP) | Planaria | 647 (v2020-03-28) | <sup>30</sup> |
| Xenopus Phenotype Ontology (XPO) | Xenopus | 20,340 (v2024-04-18) | <sup>28</sup> |
| Fission Yeast Phenotype Ontology (FYPO) | *Schizosaccharomyces pombe* | 8,056 (v2024-08-01) | <sup>8</sup> |
| Pathogen-host interaction phenotype ontology (PHIPO) | General | 1,104 (v2024-04-04) | https://github.com/PHI-base/phipo |
| Molecular glyco-phenotype ontology (MGPO) | General | 120 (v2024-04-18) | <sup>31</sup> |
| Ascomycete Phenotype Ontology (APO) | Ascomycota | 342 (v2024-04-26) | <sup>32</sup> |

In addition, phenotype standardization and integration across species is complicated by the fact that communities use different approaches to annotate phenotypes: while some use pre-composed phenotype ontology terms (for example, `HP:0007843` “Attenuation of retinal blood vessels”), others, such as Saccharomyces Genome Database (SGD) and Zebrafish Information Network (ZFIN), use a post-composed approach: the different constituents of the phenotype are captured individually during the curation process using several terms from multiple domain-specific ontologies (for example: `GO:0061304` “retinal blood vessel morphogenesis” - `PATO:0002302` “decreased process quality”)^9^.

While each approach to describing phenotypes provides standardization for the use cases of a specific community, comparing phenotype data from more than one species at scale is difficult and/or very time consuming, as it cannot be done computationally. In contrast, the use of species-neutral ontologies, such as the Gene Ontology (GO)^10^ and Uberon^11^ allow for easier interoperability across a range of taxons. GO is regularly used in many types of large-scale molecular biology experiments, including in genomics, transcriptomics, proteomics, or metabolomics. Annotations made in one species may be automatically applied to other species based on orthology, and cross-species data is easily visualized on many platforms. Similarly, Uberon, the cross-species anatomy ontology, can be used to visualize expression data across species with minimal effort^12,13^. Phenomics is a relatively young discipline and it has yet to establish a similar level of standardization to GO^14,15^. Phenomics vocabularies are developed largely as independent, siloed projects by different communities for different purposes, are often species-specific, and even within the same organism can have multiple, incompatible representations. This ultimately hampers the ability to compare phenotype data across species.

A species-neutral phenotype ontology could integrate phenotype data from any species, allowing for more straightforward incorporation of emerging model organisms and other non-model species, and expanding the scope of comparative biological research. Tools that combine human-specific data with data from only one or two model organisms have already proven significant performance gains in variant analysis^16^, disease diagnosis^17^, and potential disease model identification^18^. Such integrated data could help clinicians to select models that best address their research questions; identify cases where phenotypes are or are not associated with variants in orthologous genes; and uncover factors that influence disease penetrance and severity^19^. Phenotype integration also provides a robust approach to align molecular-level phenotypes across large evolutionary distances. For example, the reuse of phenotypic data and variant associations from yeast models such as *Saccharomyces cerevisiae* and *Schizosaccharomyces pombe* can enable predictions related to the molecular basis for diseases, particularly when conserved residues are present in human orthologs.

Efforts such as the Monarch Initiative, the Alliance of Genome Resources, Planteome, and PhenomeNET have integrated a selection of phenotype data employing a variety of methodologies such as the Entity-Quality (EQ) methodology^13,20–22^ and lexical and logical matching. The uPheno framework described here builds upon and improves these approaches to establish a unified structure for capturing phenotypic information across species, maintained as a community initiative and applied to a variety of cross-species use cases including clinical diagnostics, data discoverability, and data standardization.

## 2. Results

### 2.2. Unified Phenotype Ontology (uPheno) framework

We have developed uPheno, a framework for cross-species integrative phenomics. uPheno has three main components: the uPheno ontology; a library of design patterns and templates for computationally tractable phenotype definitions; and a number of standardized mappings to connect disparate phenotype ontologies. The uPheno ontology currently integrates 12 species-specific phenotype ontologies (Table 1), which are used by a wide range of databases from the domain of model organisms, including all databases participating in the Alliance of Genome Resources^13^, and leverages previous efforts to integrate species-specific anatomy ontologies, most notably in the Uberon ontology^23^ and Cell Ontology (CL)^24^.

Every phenotype term in the uPheno ontology represents a deviation from a reference phenotype (for example, wild-type) defined using a specific design pattern from our library (see Data Availability section). This enables phenotype classes to be defined in a consistent logical framework rather than defining each phenotype class manually. For example, the phenotype term `UPHENO:0001471` “increased size of the heart” can automatically be generated, along with labels and logical axioms, and accurately classified by instantiating a *increasedSizeOfAnatomicalEntity* pattern with a `UBERON:0000948` “heart” class from the anatomy ontology Uberon. In addition to the uPheno ontology, which includes logical connections to all species-specific ontologies, standardized mapping tables are provided with direct links between species-specific and species-neutral ontologies.

#### 2.2.1. Library of computational phenotype patterns

The majority of ontologies in biomedical sciences, especially those covering model organisms, are curated manually using tools such as Protege^25^. The use of design patterns to augment ontology development processes in the OBO (Open Biological and Biomedical Ontologies^26^) community became popular with the emergence of easy-to-use templating systems such as DOSDP^27^. Rather than manually specifying a term such as “abnormally increased glucose levels in the blood” (which includes writing a human-readable definition, a label and logical axioms), a DOSDP pattern defines a template for terms of the type “abnormally increased X levels in the Y”, including the exact structure of the label, definition and all its surrounding axioms. This ensures that all terms are consistently labeled (which is particularly difficult in large-scale ontologies such as the phenotype ontologies) and consistently axiomatized. With the addition of reasoning to ontology build pipelines, this consistent axiomatization drives consistent classification (i.e. organization in a hierarchical structure). Some ontologies, such as ZP or XPO^28^, are entirely bootstrapped from phenotype patterns (see Discussion section), which reduces the overhead of maintaining them.

We have developed 262 phenotype term templates that cover cases such as “abnormally increased X levels in the Y” (where X is a chemical entity and Y is an anatomical location), “abnormal X morphology” (where X is an anatomical entity) or “abnormal rate of X” (where X is a biological process). Details on the engineering methodology can be found in the Methods section. All patterns are available as part of a library of phenotype patterns on GitHub (see Data Availability section).

Phenotypes that affect anatomical entities (`UBERON:0001062`) or biological processes (`GO:0008150`) feature prominently in the shared uPheno pattern library constituting approximately 75% of the pattern templates (Figure 1). Patterns involving anatomical entities make up over 65% of patterns and cover both the morphology and physiology of these entities. Examples involving abnormal anatomical entities include the pattern *abnormalLengthOfAnatomicalEntity* which can be applied to phenotypes characterized by the abnormal length of any anatomical entity, such as `HP:0200011` “Abnormal length of corpus callosum”, `MP:0011999` “abnormal tail length”, and `ZP:0022039` “head length, abnormal”.

**Fig. 1.**
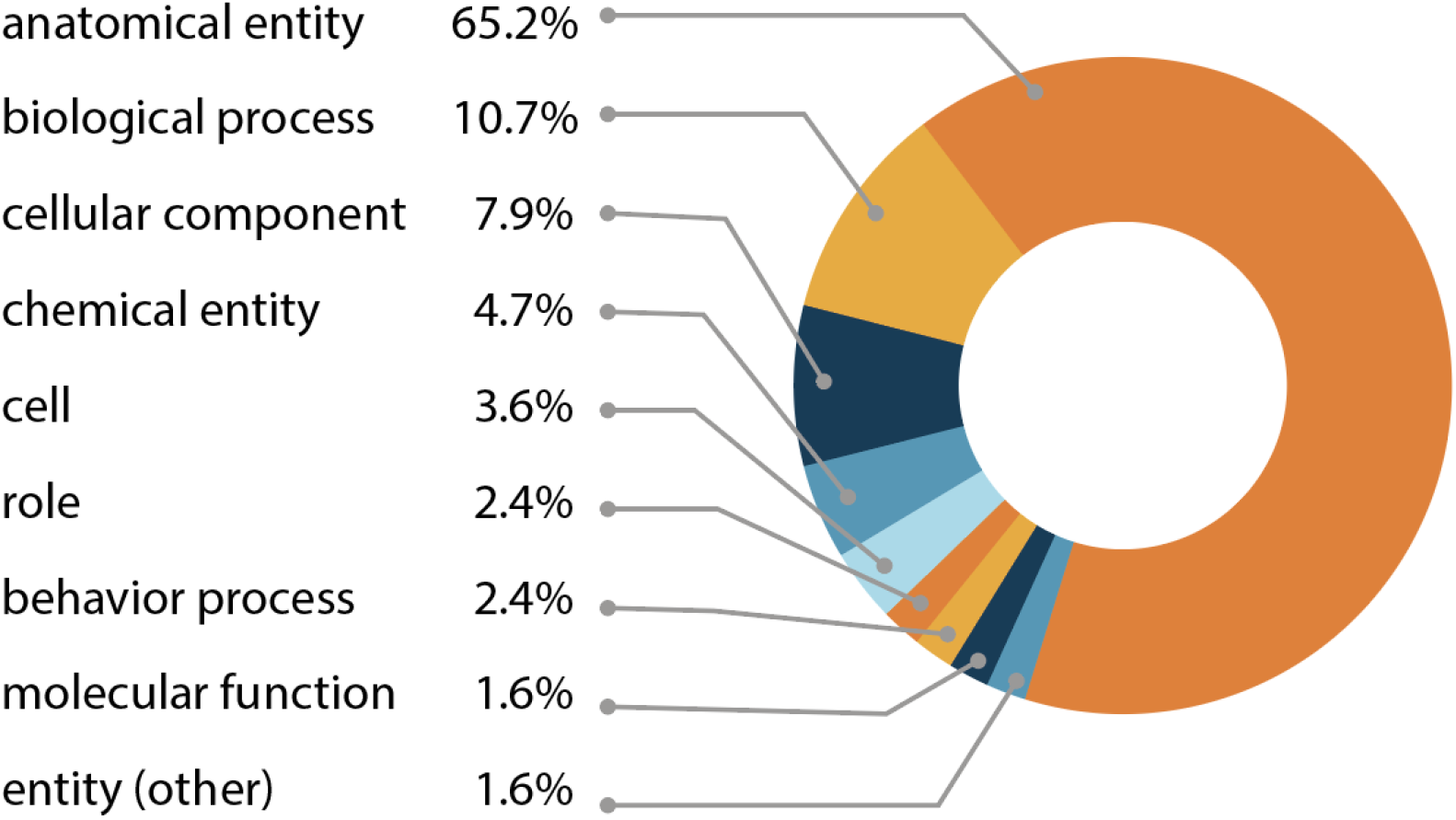
Distribution of entity types in the uPheno pattern library. All phenotype definitions reference at least one affected entity. The percentage of patterns using an entity type relative to all pattern templates are indicated. The main entity categories in uPheno phenotype pattern templates include: anatomical entity (UBERON:0001062), biological process (GO:0008150), cellular component (GO:0005575), chemical entity (CHEBI:24431), cell (CL:0000000), role (CHEBI:50906), behavior process (NBO:0000313), molecular function (GO:0003674), other entities (BFO:0000001).

Besides phenotypes described by anatomical entity abnormalities, researchers often report the alterations in biological processes associated with specific genetic mutations. The second most frequent phenotype pattern group in the uPheno library relates to biological processes (10.7%). For example, the pattern *abnormallyDecreasedRateOfContinuousBiologicalProcess* can be applied to such diverse process phenotypes as `MP:0020234` “decreased basal metabolism”, `ZP:0101378` “glycolytic process decreased rate, abnormal”, `FBcv:0000791` “decreased speed of aging”, `ZP:0001531` “blood circulation decreased rate, abnormal”, `ZP:0102933` “digestion decreased rate, abnormal”, and `FYPO:0000419` “decreased rate of cytokinesis”.

Over 11% of the pattern templates involve cellular (`CL:0000000`) or cellular component (`GO:0005575`) phenotypes. Cell component phenotypes can be observed in both single- and multicellular organisms, thus making the relevant uPheno pattern templates applicable to diverse taxa. For example, the *abnormallyDecreasedNumberOfCellularComponent* pattern template can be used in cases where a decrease in the number of mitochondria is observed, such as `HP:0040013` “Decreased mitochondrial number”, `MP:0011629` “decreased mitochondrial number”, `DDPHENO:0000271` “decreased number of mitochondria”, and `FYPO:0003820` “mitochondria present in decreased numbers during vegetative growth”.

Other templates allow the standardization of phenotypic description and annotation related to chemical entities (`CHEBI:24431`), chemical roles (`CHEBI:50906`), behavioral processes (`NBO:0000313`), molecular function (`GO:0003674`), and developmental processes (`GO:0032502`).

#### 2.2.2. uPheno ontology

The uPheno ontology is a logic (OWL) based ontology that combines existing phenotype ontologies into a single ontology and introduces common grouping classes such as `UPHENO:0082544` “mitochondrion phenotype”. For example, `HP:0001640` “Cardiomegaly”, `MP:0000274` “enlarged heart” and `ZP:0000532` “heart increased size, abnormal” all classify under a common species-neutral grouping `UPHENO:0001471` “increased size of the heart”, which is in turn classified under `UPHENO:0075162` “size of heart phenotype”. The grouping classes are primarily built using the uPheno pattern library and rely on external species-neutral ontologies for the component parts. For example, anatomical phenotype terms are created using anatomy terms from Uberon^11,23^, cell type phenotype terms from CL^24^ and physiological and subcellular phenotypes from GO^10^.

The overall structure of the uPheno ontology relies heavily on the structure of the ontologies used to build the classes. By using a well established “entity quality” (EQ) modeling framework (see Methods section), we can define a phenotype in terms of its constituent parts (for example, an anatomical and a chemical entity) and use the hierarchical structure of the respective source ontologies for these parts to classify our phenotype terms. For example, the phenotype `UPHENO:0047922` “increased thickness of the aortic valve leaflet” is classified as a “heart morphology phenotype” (`UPHENO:0076810`), because “thickness” (`PATO:0000915`) is considered a subclass of “morphology” (`PATO:0000051`) in the PATO ontology and the “aortic valve” (`UBERON:0002137`) is considered a part of the “heart” (`UBERON:0000948`). For details about this approach refer to the Methods section. There are currently 35782 terms in uPheno, of which only 7 are manually classified (all are grouping classes such as `UPHENO:3000006` “taste/olfaction phenotype” which are difficult to define using a simple EQ, see Methods).

The uPheno classes enable expressive querying and effective classification of phenotypes across species. For example, a user might want to find all genes where perturbations alter heart morphology regardless of species. To achieve this, they can simply retrieve all subclasses of “heart morphology phenotype” (`UPHENO:0076810`), for example using the Ontology Lookup Service (OLS, https://www.ebi.ac.uk/ols4/ontologies/upheno) (Fig. 2(C)). This straightforward retrieval of similar phenotypes across taxa can be used for a variety of applications, such as finding relevant literature across species, identifying candidate genes for phenotypes with an unknown genetic basis, or comparing the phenotypic spectrum produced by mutations in orthologous genes across species. The use of the EQ logical framework moreover enables querying for phenotypes using logical expressions. For example, a bioinformatician interested in heart morphology phenotypes across species could use an OWL class expression `(‘has part’ some (morphology and (‘characteristic of part of’ some (‘heart’)) and (qualifier some abnormal))`.

**Fig. 2:**
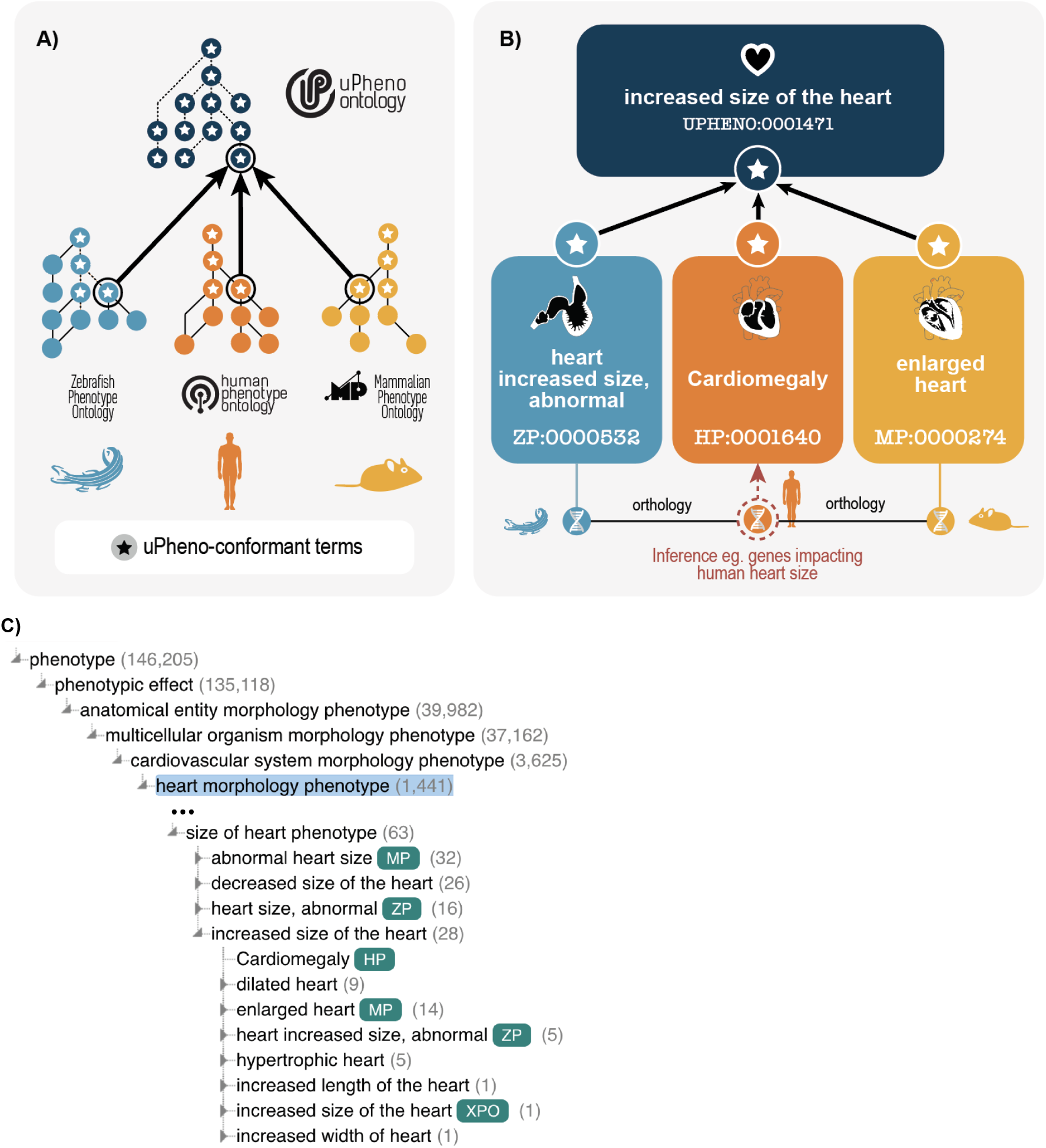
A. uPheno is a framework for consistent and logical definition of phenotype categories using ontology design patterns that provides a hierarchical vocabulary of species-neutral phenotype terms under which their species-specific counterparts are grouped. The ontology design templates are based on shared features of existing phenotypic descriptions from various model organisms and represent community consensus. The phenotype-pattern template adherent terms are adopted by species-specific ontologies, thereby contributing to the community-built uPheno framework. B. uPheno accelerates cross-species inference and computationally amenable comparative phenotype analysis. For example, the interoperable representation of heart phenotypes characterized by increased size, compared with wild-type in distinct species, such as zebrafish and human, allows the cross-species identification of genes whose alleles can cause similar phenotypes. C. uPheno contextual hierarchy for increased size of the heart.

uPheno currently integrates 12 species-specific phenotype ontologies to varying degrees; see Table 1. The deepest level of integration is for ontologies that cover vertebrates, such as HPO, MP, ZP, and XPO. The integration of other ontologies is more variable; for example, WBPhenotype, DDPHENO and DPO are well integrated, while FYPO and APO are at earlier stages of integration.

For curation scenarios where no species-specific vocabulary exists, uPheno provides a standardized species-neutral vocabulary that can be used to capture phenotype data (see the Online Mendelian Inheritance in Animals (OMIA) example in Discussion Section).

#### 2.2.3. Cross-species mappings

Cross-species mappings can be used to make datasets interoperable across species, for example by linking HP phenotypes from a human study to similar MP phenotypes in a mouse study. To facilitate these types of integrations we publish a number of cross-species mappings derived from the cross-species ontologies (e.g Uberon, GO) used in the logical definitions of the terms. For example, `MP:0003855` “abnormal forelimb zeugopod morphology” maps to `HP:0002973` “Abnormal forearm morphology” as they both are defined using the same anatomical term `UBERON:0002386` “forelimb zeugopod”.

These mappings are published in a simple, standardized spreadsheet using the SSSOM^33^ TSV standard, connecting phenotype terms from one species-specific phenotype ontology such as ZP to another, such as XPO. uPheno semantic similarity tables provide associations between species-specific phenotype terms and scores that reflect their semantic similarity. All cross-species mappings, semantic similarity tables and manually curated mappings can be obtained from the URLs provided in the Data Availability Section.

## 3. Methods

uPheno integrates existing species-specific representations of phenotype data developed by a broad community of model organism, clinical, and research database curators, using a variety of methodologies. Many of these representations already exist as pre-composed (also known as pre-coordinated) ontologies, where specific terms such as “decreased circulating lysine level” are created and assigned unique, permanent identifiers (“MP:0030719”). Other representations instead rely on curating the different aspects of phenotype separately (post-composition, also known as post-coordination). For example, ZFIN^34^ follows a sophisticated post-composed curation style, selecting the attribute and the entity terms separately. To facilitate the integration of this phenotypic data, the ZP ontology, supported by the uPheno effort, converts the post-composed curated content in the ZFIN database into a pre-coordinated ontology.

### 3.1. EQ framework

The computational phenotype model underlying the uPheno framework is an extension of the entity–attribute (or Entity-Quality, EQ) model which is used to describe phenotypes in terms of affected entities and their characteristics^35^. The affected entities included in phenotypic characterizations are called the *bearers* of the observable attributes (also known as observable qualities or characteristics). For example, in the phenotype ‘enlarged heart’ the entity, heart, bears the characteristic or quality of increased size.

The attribute categories (qualities) in uPheno logical axioms that characterize phenotypes are chosen from the Phenotype and Trait Ontology (PATO^36^). The entities in uPheno can be broadly categorized as physical objects or processes. The physical objects include anatomical entities and their constituents, such as cells, subcellular structures or components, proteins and other chemical entities. Examples of process-type entities include GO biological process and GO molecular function classes as well categories of behavior or roles for chemical compounds. The entity components in uPheno include classes from OBO Foundry ^26^ ontologies, such as Uberon, the Cell Ontology, GO (cellular component classes), the Chemical Entities of Biological Interest (ChEBI), and the Neuro Behavior Ontology (NBO)^10,23,24,37,38^.

The most basic EQ model involves two primitive classes that are part of an asserted class expression, where the “E” entity is the affected entity (e.g. an anatomical entity, such as “limb”) or a biological process (e.g. “limb development”). The “Q” component is a quality (attribute) class from PATO. The equivalent class axioms in uPheno follow or extend the basic EQ model to represent phenotypic characteristics. uPheno is intended to represent phenotypic states that deviate from a reference, therefore they always include a `PATO:0000460` “abnormal” component. This is expressed as an equivalent class axiom. For example `UPHENO:0076810` “heart morphology phenotype” is expressed in OWL Manchester syntax^39^ as follows:

~~~
has part some (
  morphology and
  characteristic of part of some heart and
  has modifier some abnormal)
~~~

The relationships `RO:0000052` “characteristic of” and `RO:0002314` “characteristic of part of” from the OBO Relations Ontology (RO) ^40^ are used to connect the entity that is observed to be phenotypically affected and the characteristic it exhibits (e.g. “size” or “amount”). The entity part of the EQ statement can be composite, i.e. comprising more than one entity. For example, if an abnormality of a biological process occurs in a particular anatomical location, then the entity will be defined as a biological process (`GO:0008150`) which “occurs in” (`BFO:0000066`) an “anatomical location” (`UBERON:0001062`). More complex patterns defining entities can be found, such as chemical entities that play a certain role (for example a `CHEBI:25212` “metabolite” in an `UBERON:0001062` “anatomical location”).

This formal logic representation of phenotypes in OWL enables the use of logical inference through automated reasoners; see Section on the uPheno Ontology above for an example on how the use of EQ statements in conjunction with reference ontologies such as ChEBI and Uberon enables entirely automated classification of phenotypes. More information about how OWL ontologies and reasoning can be leveraged in the biological and biomedical sciences can be found in Hoehndorf et al^41^.

### 3.2. The DOSDP framework (and ODK)

Dead Simple OWL Design Patterns (DOSDPs)^27^ allow efficient and scalable definition of ontology term templates which can support the construction and maintenance of large numbers of ontology classes. The EQ modeling framework is especially well suited for template-based ontology development. DOSDP term templates, which are specified in YAML, support specification of both logical axioms and annotation axioms (e.g. for synonyms, labels and definitions) with variable slots. Separate tables (stored as tsv or csv files) specify fillers for these variables. The templates and tables can be parsed and converted into OWL axioms by dosdp-tools, which is part of the Ontology Development Kit (ODK)^42^. The resulting axioms can be built into a class hierarchy and/or incorporated into an existing OWL ontology by ROBOT^43^ and other ODK components. The release system of the uPheno ontology is implemented as an ODK workflow, which makes it easily executable in a platform-agnostic manner through Docker.

### 3.3. uPheno templates

uPheno phenotype pattern templates are designed to help align the modeling of similar or related phenotypic categories across multiple taxonomic domains. uPheno utilizes the DOSDP templating system and the EQ framework to define phenotype templates. The uPheno templates are the result of collective curation by a community of ontology editors called the Phenotype Ontology Reconciliation Effort^44^. This community effort is pivotal not only to the definition of phenotype templates described in this section, but also in their implementation in the species-specific phenotype ontologies. The curation process operates as follows: when a need for a pattern arises, a member of the community requests a template. Next, another member of the community develops a draft template in DOSDP format and makes a pull request on GitHub. The broader community can review the pattern and provide feedback suggesting changes to wording, definitions, naming templates and other aspects. Once the template is approved, it is presented at a specific monthly call that is organized by the reconciliation effort for the purpose of advancing uPheno patternization across the phenotype ontology editors community. All present members who approve of the pattern now add their signature to the template (in the form of an ORCID, see below), indicating their approval and their intention to implement the pattern in the phenotype ontology they represent (for example, HPO, MP, DDPHENO, etc).

As an example, a slightly simplified version of the *abnormalAnatomicalEntity* pattern can be seen in Figure 3. The full pattern can be downloaded at the URL indicated by the “pattern_iri” field.

**Figure 3:**
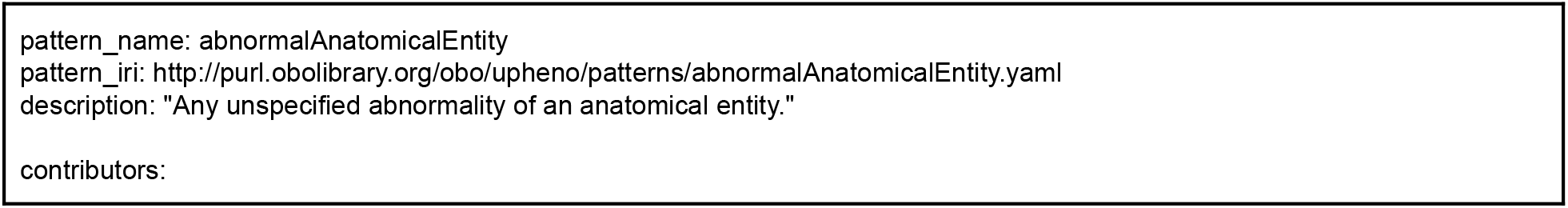
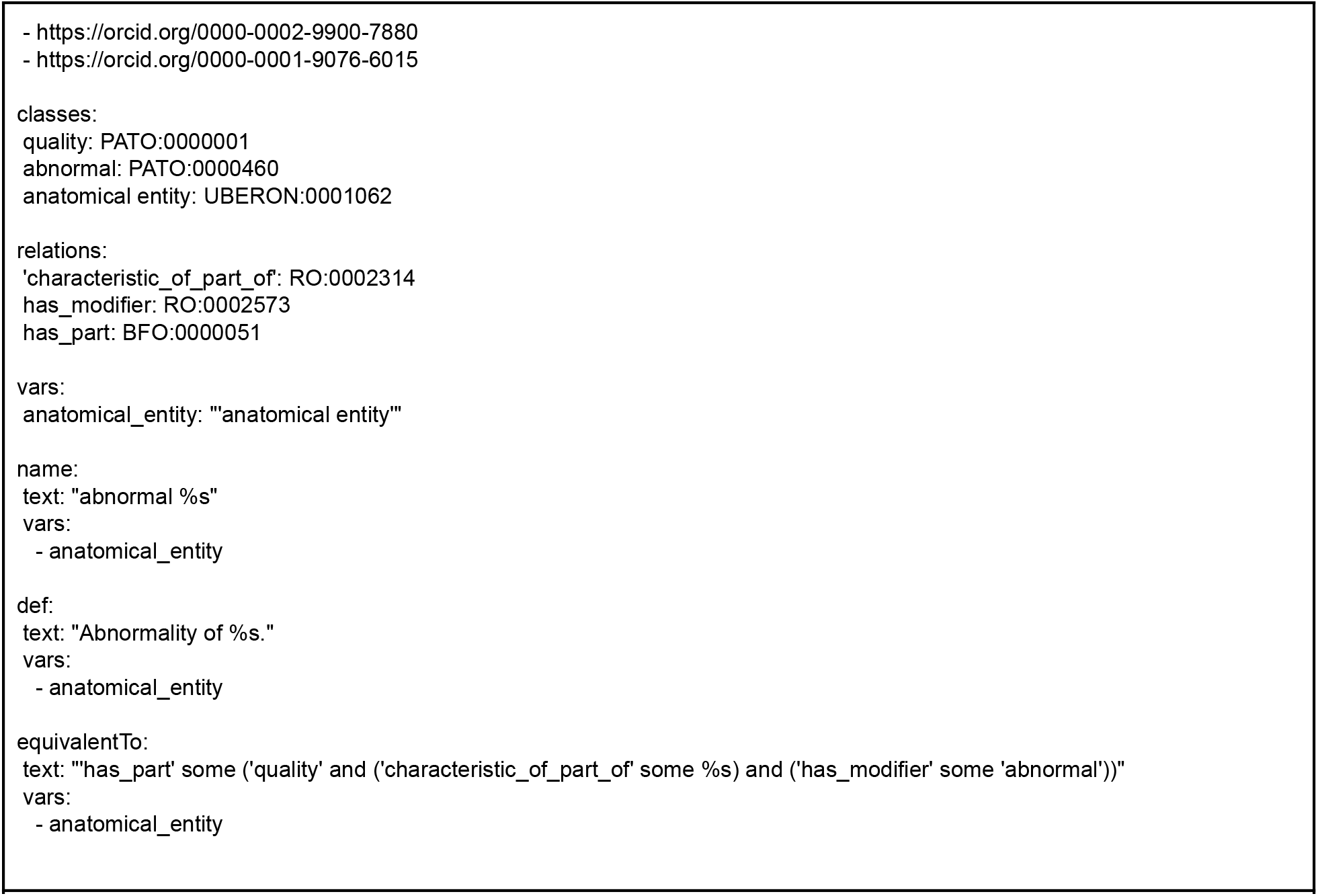
DOSDP pattern for the representation of abnormal anatomical entity phenotypes. Species-specific phenotype ontologies implement this pattern in phenotype terms such as “Abnormality of the cardiovascular system” (HP:0001626) and “gall bladder quality, abnormal” (ZP:0006529).

The two most important elements of any pattern are the template for the logical definition (`equivalentTo`) and the list of contributors. The `equivalentTo` field specifies the logical axiom pattern to be used to define the phenotype term. The `contributor` field is used to record the members of the reconciliation effort who have reviewed a particular pattern (see above).

The main relationships used by phenotype patterns can be seen in Table 2. Other relationships are used in specific cases. For example, `GOREL:0001006` “acts on population of” is used for the definition of cell proliferation patterns in order to align with the logical definition of cell proliferation process terms in GO.

**Table 2:** Relationships used to logically define terms in uPheno.

| Relationship | Meaning |
| --- | --- |
| characteristic of | Relates the phenotypic quality to the “bearer”, i.e. the entity that is affected by the phenotype. |
| characteristic of part of | Relates the phenotypic quality to the “bearer” or one of its parts. |
| part of | A general mereological relation that denotes parthood between two entities, such as an anatomical entity and one of its parts. |
| has part | The inverse relationship of part of. In the context of our EQs, this is used to relate the phenotype itself to its primary quality. |
| towards | In the case of a relational quality, i.e. a quality that involves two entities (such as “fused with”), the towards relation is used to describe the second involved entity (the first is related using “characteristic of”). |
| has modifier | Phenotypes can be either normal or abnormal. The relation is used to connect the primary phenotypic quality to a respective modifier. |
| occurs in | Used to connect biological processes to the anatomical entities in which they are taking place. |
| Table 2: Relationships used to logically define terms in uPheno |  |

The uPheno patterns described here are collected and available in the “uPheno pattern library”, which makes it easy for phenotype ontology editors to identify a suitable pattern for a given phenotype (see Data Availability).

### 3.4. Cross-species mappings and Semantic similarity

Locating genetic variants with similar phenotypes across species can suggest new disease candidates, or provide new insights into gene function. For example, *PRG4* mutations have been causally implicated in “Camptodactyly of finger” (`HP:0100490`) phenotypes in human genetic studies, and there are mouse knock-out models of the orthologous *Prg4* gene whose phenotype annotations include “camptodactyly” (`MP:0003807`), thereby providing independent evidence of causality and increasing confidence that the gene-to-phenotype association is correct^5,45,46^. To leverage these similar phenotypes for data analysis (especially when potentially orthologous variants relating to analogous phenotypes are unknown), we publish those mappings in a standardized format called the Simple Standard for Sharing Ontological Mappings (SSSOM^33^). SSSOM allows the precise specification of the mapping relation, such as `semapv:crossSpeciesExactMatch`, to signify a relation between two identical phenotypic characteristics of homologous anatomical structures. Metadata can also be attached to distinguish matches which have been determined through manual curation (`semapv:ManualMappingCuration`), lexical matching (`semapv:LexicalMatchin`g) and logical matching using automated reasoning (`semapv:LogicalMatching`). Mapping tables are particularly useful for basic lookup tasks and providing cross-links between resources. For example, IMPC ^47^ and MGI ^48^ websites use mapping tables to allow searching using an HPO phenotype name to discover data connected to an analogous MP phenotype term.

The uPheno framework also enables the computational identification of semantically similar phenotypes. Similar phenotypes are more likely to be related to similar mechanisms, which makes this information very valuable for applications such as variant prioritization. For example, Huntington’s disease and Parkinson disease are both neurodegenerative disorders where involuntary motor symptoms, chorea and tremors respectively, are prominent phenotypic features. *HTT* gene mutations have been causally implicated in Huntington’s disease. We could therefore predict *HTT* might be a candidate gene for Parkinson disease based on phenotypic similarity. This is exploited to prioritize pathogenic variants in tools such as Exomiser^16^, which is used widely in clinical practice. Estimating “phenotypic similarity” in Exomiser uses two main measures: Jaccard, which is simply a measure of how similar two phenotypes are with respect to their position in the ontology (sibling terms are more similar than distantly related terms); and the PhenoDigm score^49^, which is based on Jaccard similarity but normalizes against Information Content, which itself is a measure of “how informative/interesting” a phenotype is (phenotypes that are more specific, have stronger known gene associations, and are positioned lower in the ontology hierarchy are considered “more informative”).

## 4. Discussion

### 4.1. uPheno applications

uPheno is applicable to a variety of phenotypic analysis projects and tools. The International Mouse Phenotyping Consortium (IMPC)^47^, a global resource for whole gene knockout (KO) mouse lines, plans to use uPheno cross-species mappings to make mouse phenotypes discoverable using HPO phenotype terms.^47^ Similarly, MGI has recently prototyped the use of cross-species mappings for discovering gene-to-phenotype associations^48^ and intends to incorporate uPheno mappings into this tool. The Monarch Initiative Knowledge Graph^20^ uses uPheno alongside the Ontology of Biological Attributes (OBA),^50^ which enables analyzing biomedical data across species, with a specific focus on phenotypes, diseases, and their genetic underpinnings.

uPheno has also been applied to the phenomics informed study of disease. In a study to determine whether model organism phenotype data contributes to the computational discovery of human gene-disease associations and to what extent, Alghamdi et. al. used uPheno and Pheno-e^51^ (an extension of the PhenomeNET ontology), to semantically relate phenotypes resulting from loss-of-function mutations in mouse, zebrafish, fruit fly and fission yeast model organisms, to disease-associated human phenotypes^52^. An informatics pipeline developed by Cary et al. presented an Alzheimer’s disease risk assessment score across biological domains. The approach utilized phenotypes of model organism orthologs to human genes extracted from the uPheno ontology^53^. InpherNet is a machine-learning approach that can aid monogenic disease diagnosis where patient-based annotation is incomplete or lacking. It leverages the uPheno ontology to obtain organismal and cellular level gene phenotype data.^54^

In disease diagnostics, variant prioritization tools such as Exomiser^16^, EmbedPVP^55^, and EvORanker^56^ leverage uPheno’s cross-species phenotypic similarity mappings to improve variant prioritization by comparing human phenotypes to those of model organisms like mice and zebrafish. The Phenotypic Inference Evaluation Framework (PhEval)^57^ has recently been developed to benchmark diagnostic yield in Exomiser when informed by similar cross-species phenotypes mapped using uPheno.

uPheno can also be used to bootstrap the generation of species-specific phenotype ontologies. Instead of building an ontology manually, uPheno pattern templates and spreadsheets of relevant entities can be used to automate the creation of ontology terms. Xenbase^58^, the Xenopus model organism knowledgebase, has developed the Xenopus phenotype ontology (XPO^28^) using uPheno templates in combination with high-level terms from the Xenopus Anatomy Ontology (XAO^59^), the Phenotype and Trait Ontology (PATO) and the Gene Ontology (GO). The PLANP ontology for the Planarian Flatworm has been generated using uPheno patterns for use in phenotype annotation. These terms are being used to help researchers identify genes with comparable phenotypes when perturbed using RNA interference.

### 4.2. Limitations

While uPheno allows the species-neutral description of a phenotype such as “abnormally enlarged heart”, it does not address what reference the phenotype is a comparison to (e.g. a control group or wild type), nor does it capture an effect size (e.g. whether the phenotype is slightly outside of the clinically normal range or significantly changed). In practice all phenotype ontologies are used in contexts where different comparators and effect sizes are assumed. For example, for quantitative traits in the GWAS Catalog, the annotation with a phenotype/trait term indicates that the effect allele is associated with an increase/decrease in the trait compared to the mean of the entire sample; for binary traits, the trait annotation indicates that the effect allele is found at higher/lower frequency in cases compared to controls. Since the overall goal of uPheno is to make phenotype information comparable, it would be impractical to create different classification axes for every case (e.g. having an “abnormally increased heart size”, a “significantly increased heart size”, a “abnormally increased heart size compared to wild-type”). Thus, the presence of a phenotype term as part of, for example, a gene-to-phenotype association, cannot automatically be associated with a specific comparator or effect size. Instead, this information needs to be supplied in the metadata of the phenotype annotation, for example, the experimental conditions.

A second important limitation is that, while the uPheno framework has significantly improved the alignment of species-specific phenotype ontologies with each other and with uPheno, it does not automatically lead to their complete alignment. The implementation of uPheno patterns with uPheno-conformant reference ontologies such as Uberon or Uberon-aligned ontologies by the species-specific phenotype ontologies is costly in developer time, so coverage for ontologies such as HPO and MP is unlikely to be complete in the near future. This is further aggravated by the existence of complex phenotype classes which cannot be defined logically using uPheno patterns such as “hemifacial hypoplasia” (`HP:0011332`, `MP:0030100`) or “anencephaly” (`HP:0002323`, `MP:0001890`). Nevertheless, significant coverage has already been reached, and the process of alignment is ongoing.

### 4.3. Future work

In the future, we would like to make uPheno more accessible to researchers by integrating the ontology into tools to enable use cases such as finding related phenotypes across species without the need for specialized ontology training. We also plan to improve the upper level structure of the phenotype hierarchy to improve findability of phenotype terms for users browsing the ontology.

uPheno has primarily focused on integrating phenomics data from the model organism community. We are expanding our efforts to include non-model animal species and address use cases relevant to the veterinary field. This undertaking has been driven by the team at the Online Mendelian Inheritance in Animals (OMIA, https://omia.org/home/) ^60^ with whom we are collaborating. OMIA is a freely available, curated knowledge base that offers up-to-date information on inherited disorders, traits, and associated genes and variants in animals. uPheno was chosen to represent phenotypes, clinical and pathological signs data in OMIA to enhance their data’s computational analysis and data interoperability.

## 5. Data Availability Statement

The uPheno ontology, pattern library and associated files can be found here: https://github.com/obophenotype/upheno/blob/master/docs/reference/data-availability.md, or at the URLs provided in the text. The uPheno ontology can be browsed using OLS (https://www.ebi.ac.uk/ols4/ontologies/upheno).

## Literature cited

1. Lima Cunha, D., Arno, G., Corton, M. & Moosajee, M. The Spectrum of PAX6 Mutations and Genotype-Phenotype Correlations in the Eye. Genes 10, (2019).

2. Fisher, S. E. & Scharff, C. FOXP2 as a molecular window into speech and language. Trends Genet. 25, 166–177 (2009).

3. Rodgers, B. D. & Garikipati, D. K. Clinical, agricultural, and evolutionary biology of myostatin: a comparative review. Endocr. Rev. 29, 513–534 (2008).

4. Gargano, M. A. et al. The Human Phenotype Ontology in 2024: phenotypes around the world. Nucleic Acids Res. 52, D1333–D1346 (2024).

5. Smith, C. L. & Eppig, J. T. The mammalian phenotype ontology: enabling robust annotation and comparative analysis. Wiley Interdiscip. Rev. Syst. Biol. Med. 1, 390–399 (2009).

6. Osumi-Sutherland, D. et al. The Drosophila phenotype ontology. J. Biomed. Semantics 4, 30 (2013).

7. Schindelman, G., Fernandes, J. S., Bastiani, C. A., Yook, K. & Sternberg, P. W. Worm Phenotype Ontology: integrating phenotype data within and beyond the C. elegans community. BMC Bioinformatics 12, 32 (2011).

8. Harris, M. A., Lock, A., Bähler, J., Oliver, S. G. & Wood, V. FYPO: the fission yeast phenotype ontology. Bioinformatics 29, 1671–1678 (2013).

9. Mungall, C. J. et al. Integrating phenotype ontologies across multiple species. Genome Biol. 11, R2 (2010).

10. Ashburner, M. et al. Gene ontology: tool for the unification of biology. The Gene Ontology Consortium. Nat. Genet. 25, 25–29 (2000).

11. Mungall, C. J., Torniai, C., Gkoutos, G. V., Lewis, S. E. & Haendel, M. A. Uberon, an integrative multi-species anatomy ontology. Genome Biol. 13, R5 (2012).

12. Gene Ontology Consortium et al. The Gene Ontology knowledgebase in 2023. Genetics 224, (2023).

13. Bult, C. J. & Sternberg, P. W. The alliance of genome resources: transforming comparative genomics. Mamm. Genome 34, 531–544 (2023).

14. Brown, S. D. M. et al. High-throughput mouse phenomics for characterizing mammalian gene function. Nat. Rev. Genet. 19, 357–370 (2018).

15. Rahman, J. & Rahman, S. The utility of phenomics in diagnosis of inherited metabolic disorders. Clin. Med. 19, 30–36 (2019).

16. Smedley, D. et al. Next-generation diagnostics and disease-gene discovery with the Exomiser. Nat. Protoc. 10, 2004–2015 (2015).

17. Sun, Y. V. & Hu, Y.-J. Integrative Analysis of Multi-omics Data for Discovery and Functional Studies of Complex Human Diseases. Adv. Genet. 93, 147–190 (2016).

18. Dickinson, M. E. et al. High-throughput discovery of novel developmental phenotypes. Nature 537, 508–514 (2016).

19. Cirincione, A. G., Clark, K. L. & Kann, M. G. Pathway networks generated from human disease phenome. BMC Med. Genomics 11, 75 (2018).

20. Putman, T. E. et al. The Monarch Initiative in 2024: an analytic platform integrating phenotypes, genes and diseases across species. Nucleic Acids Res. 52, D938–D949 (2024).

21. Rodríguez-García, M.Á., Gkoutos, G. V., Schofield, P. N. & Hoehndorf, R. Integrating phenotype ontologies with PhenomeNET. J. Biomed. Semantics 8, 58 (2017).

22. Cooper, L., Elser, J., Laporte, M.-A., Arnaud, E. & Jaiswal, P. Planteome 2024 Update: Reference Ontologies and Knowledgebase for Plant Biology. Nucleic Acids Res. 52, D1548–D1555 (2024).

23. Haendel, M. A. et al. Unification of multi-species vertebrate anatomy ontologies for comparative biology in Uberon. J. Biomed. Semantics 5, 21 (2014).

24. Diehl, A. D. et al. The Cell Ontology 2016: enhanced content, modularization, and ontology interoperability. J. Biomed. Semantics 7, 44 (2016).

25. Musen, M. A. & Protégé Team. The Protégé Project: A Look Back and a Look Forward. AI Matters 1, 4–12 (2015).

26. Jackson, R. et al. OBO Foundry in 2021: operationalizing open data principles to evaluate ontologies. Database 2021, (2021).

27. Osumi-Sutherland, D., Courtot, M., Balhoff, J. P. & Mungall, C. Dead simple OWL design patterns. J. Biomed. Semantics 8, 18 (2017).

28. Fisher, M. E. et al. The Xenopus phenotype ontology: bridging model organism phenotype data to human health and development. BMC Bioinformatics 23, 99 (2022).

29. Fey, P., Dodson, R. J., Basu, S., Hartline, E. C. & Chisholm, R. L. dictyBase and the Dicty Stock Center (version 2.0) - a progress report. Int. J. Dev. Biol. 63, 563–572 (2019).

30. Nowotarski, S. H. et al. Planarian Anatomy Ontology: a resource to connect data within and across experimental platforms. Development 148, (2021).

31. Gourdine, J.-P. F. et al. Representing glycophenotypes: semantic unification of glycobiology resources for disease discovery. Database 2019, (2019).

32. Engel, S. R. et al. Saccharomyces Genome Database provides mutant phenotype data. Nucleic Acids Res. 38, D433–6 (2010).

33. Matentzoglu, N. et al. A Simple Standard for Sharing Ontological Mappings (SSSOM). Database 2022, (2022).

34. Bradford, Y. M. et al. Zebrafish information network, the knowledgebase for Danio rerio research. Genetics 220, (2022).

35. Washington, N. L. et al. Linking human diseases to animal models using ontology-based phenotype annotation. PLoS Biol. 7, e1000247 (2009).

36. Gkoutos, G. V., Green, E. C. J., Mallon, A.-M., Hancock, J. M. & Davidson, D. Using ontologies to describe mouse phenotypes. Genome Biol. 6, R8 (2005).

37. Gkoutos, G. V., Schofield, P. N. & Hoehndorf, R. The neurobehavior ontology: an ontology for annotation and integration of behavior and behavioral phenotypes. Int. Rev. Neurobiol. 103, 69–87 (2012).

38. Hastings, J. et al. ChEBI in 2016: Improved services and an expanding collection of metabolites. Nucleic Acids Res. 44, D1214–9 (2016).

39. OWL 2 web ontology language Manchester syntax (second edition). http://www.w3.org/TR/owl2-manchester-syntax/.

40. Mungall, C. et al. Oborel/obo-Relations: 2023-08-18 Release. (Zenodo, 2023). doi:10.5281/ZENODO.8263469.

41. Hoehndorf, R., Schofield, P. N. & Gkoutos, G. V. The role of ontologies in biological and biomedical research: a functional perspective. Brief. Bioinform. 16, 1069–1080 (2015).

42. Matentzoglu, N. et al. Ontology Development Kit: a toolkit for building, maintaining and standardizing biomedical ontologies. Database 2022, (2022).

43. Jackson, R. C. et al. ROBOT: A Tool for Automating Ontology Workflows. BMC Bioinformatics 20, 407 (2019).

44. The phenotype reconciliation effort - unified phenotype ontology. https://obophenotype.github.io/upheno/reference/reconciliation-effort/.

45. Marcelino, J. et al. CACP, encoding a secreted proteoglycan, is mutated in camptodactyly-arthropathy-coxa vara-pericarditis syndrome. Nat. Genet. 23, 319–322 (1999).

46. Rhee, D. K. et al. The secreted glycoprotein lubricin protects cartilage surfaces and inhibits synovial cell overgrowth. J. Clin. Invest. 115, 622–631 (2005).

47. Groza, T. et al. The International Mouse Phenotyping Consortium: comprehensive knockout phenotyping underpinning the study of human disease. Nucleic Acids Res. 51, D1038–D1045 (2023).

48. Baldarelli, R. M. et al. Mouse Genome Informatics: an integrated knowledgebase system for the laboratory mouse. Genetics 227, (2024).

49. Smedley, D. et al. PhenoDigm: analyzing curated annotations to associate animal models with human diseases. Database 2013, bat025 (2013).

50. Stefancsik, R. et al. The Ontology of Biological Attributes (OBA)-computational traits for the life sciences. Mamm. Genome 34, 364–378 (2023).

51. Hoehndorf, R., Schofield, P. N. & Gkoutos, G. V. PhenomeNET: a whole-phenome approach to disease gene discovery. Nucleic Acids Res. 39, e119 (2011).

52. Alghamdi, S. M., Schofield, P. N. & Hoehndorf, R. Contribution of model organism phenotypes to the computational identification of human disease genes. Dis. Model. Mech. 15, (2022).

53. Cary, G. A. et al. Genetic and multi-omic risk assessment of Alzheimer’s disease implicates core associated biological domains. Alzheimers. Dement. 10, e12461 (2024).

54. Yoo, B., Birgmeier, J., Bernstein, J. A. & Bejerano, G. InpherNet accelerates monogenic disease diagnosis using patients’ candidate genes’ neighbors. Genet. Med. 23, 1984–1992 (2021).

55. Althagafi, A., Zhapa-Camacho, F. & Hoehndorf, R. Prioritizing genomic variants through neuro-symbolic, knowledge-enhanced learning. Bioinformatics 40, (2024).

56. Canavati, C. et al. Using multi-scale genomics to associate poorly annotated genes with rare diseases. Genome Med. 16, 4 (2024).

57. Bridges, Y. et al. Towards a standard benchmark for variant and gene prioritisation algorithms: PhEval - Phenotypic inference Evaluation framework. bioRxiv (2024) doi:10.1101/2024.06.13.598672.

58. Fisher, M. et al. Xenbase: key features and resources of the Xenopus model organism knowledgebase. Genetics 224, (2023).

59. Segerdell, E., Bowes, J. B., Pollet, N. & Vize, P. D. An ontology for Xenopus anatomy and development. BMC Dev. Biol. 8, 92 (2008).

60. Nicholas, F. W. Online Mendelian Inheritance in Animals (OMIA): a record of advances in animal genetics, freely available on the Internet for 25 years. Anim. Genet. 52, 3–9 (2021).

